# Assay coverage reshapes metabolic diagnosis and the interpretation of pathway disturbance

**DOI:** 10.64898/2026.10.03.756464

**Authors:** Ivan Smirnov, Clara Fontaine

**Author notes:** These authors contributed equally.

## Abstract

Metabolic profiles are used to quantify disturbance, classify disease and infer biochemical function, yet these endpoints retain different information. We combine secondary analysis of 386 metabolites in six *Arabidopsis thaliana* mutation accumulation lines, known-disease classification in 129 public human metabolic-disease profiles, and an analytically solvable kinetic model. A conditional occupancy model separates annotation-bin breadth from hit count and measured bin sizes: the plant low-versus-high fitness comparison changes from a two-sided exact permutation *p* = 0.10 to *p* = 0.30 after calibration. Unsigned metabolite profiles achieve mean macro recall of 0.911 and 0.895 in separate EDTA and heparin subsets, compared with 0.624 and 0.382 for a burden baseline. However, controlled loss of 50% of held-out measurements reverses representation ranking in heparin, giving signed profiles a recall advantage of 0.108 with a conditional bootstrap interval of 0.063–0.155. A spectral decomposition identifies the within-pathway contrasts attenuated by smoothing. The kinetic model then shows that lower concentration and faster relaxation can coexist with reduced desired product flux, while a null-space criterion states when observations identify a metabolic task. The results connect assay coverage, diagnostic information and functional identifiability. Human performance is internally validated, and the kinetic predictions remain uncalibrated to patients.

## 1. Introduction

A metabolic profile is a projection of a biochemical system. It measures selected molecular pools, at selected times, under conditions shaped by genotype, environment, sample processing and treatment. The same profile may be used to ask three substantially different questions: how broadly metabolism has changed, which disease resembles the observed pattern, and whether essential biochemical function has deteriorated. An analysis that answers one question need not answer the others.

Untargeted metabolic profiling has extended laboratory screening of inherited metabolic disorders beyond predefined analyte panels (Miller et al., 2015; Coene et al., 2018). Studies of urea-cycle disorders further show that profiles contain disease-associated alterations alongside signals reflecting dietary and medical management (Burrage et al., 2019). This motivates a distinction between a recognizable disease pattern, the completeness of its measurement, and the biochemical function it represents.

Kreutzmann et al. (2022) quantified metabolic changes in *A. thaliana* mutation accumulation lines with contrasting field-fitness rankings. Their association between lower fitness and broader metabolic disturbance provides an empirical foundation for asking how mutational effects propagate through metabolism. The published supplementary table makes a coverage-conditioned audit possible, while its aggregate format limits causal interpretation. In human disease, Thistlethwaite et al. (2022) established that network-organized patterns in untargeted metabolomics can support identification of inherited metabolic disorders. Gao et al. (2025) subsequently investigated diagnostic signatures in small and evolving rare-disease datasets. These studies motivate a clinically relevant question: what information survives when a diagnostic assay observes only part of a metabolic pattern?

Existing methods address several forms of incomplete observation. Mummichog uses network organization to infer activity from spectral features before exhaustive compound identification (Li et al., 2013); PIUMet integrates untargeted metabolite features with molecular networks (Pirhaji et al., 2016); and metPropagate uses metabolomic alterations and functional interaction networks for disease-gene prioritization (Graham Linck et al., 2020). These approaches establish useful network context but solve different inference problems.

Contemporary network methods can distribute statistical signals across partially observed reaction graphs (Kuligowski et al., 2026). Such distribution is a representation operation, not a measurement of missing concentrations. Direction-aware integration is also established: Reafect combines metabolite magnitude and sign with reaction distance and directionality for gene prioritization (Bongaerts et al., 2022). The relevant question here is how the diagnostic value of this information changes with assay coverage. Similarly, mechanistic models can link biochemical constraints to growth-related phenotypes (Wendering et al., 2025), but biochemical snapshots alone may not identify kinetic control parameters (Shin et al., 2026). Metabolic control theory distinguishes the control of concentration from the control of flux (Kacser and Burns, 1995; Heinrich and Rapoport, 1974), while stability analysis characterizes the response of metabolic states (Grimbs et al., 2007). Structural and practical identifiability provide a complementary framework for determining what a partially observed dynamical system can reveal (Raue et al., 2009; Villaverde et al., 2016). The unresolved issue addressed here is how coverage, information compression and functional observability interact in concrete empirical analyses.

We investigate three linked hypotheses without assuming that network smoothing or signed data must outperform simple alternatives. First, raw pathway breadth can partly reflect sampling combinatorics. Second, the relative value of signed and unsigned diagnostic profiles can change with measurement completeness. Third, improvement in an observed metabolic state need not certify improvement in a desired flux. We test the first two using public experimental data and establish the third using an explicit kinetic counterexample and a local identifiability criterion. We do not pool plant and human observations, transfer a plant-trained predictor to patients, or equate plant fitness with human clinical severity.

## 2. Results

### 2.1. Conditional occupancy changes the interpretation of pathway breadth

The plant supplementary workbook contains 386 metabolite fold changes and per-line *p* values for three low-fitness lines (L44, L73 and L75) and three high-fitness lines (L53, L61 and L119). We retain the 59 annotation labels present in the workbook, including the partially characterized category. These are annotation bins rather than 59 experimentally verified biochemical pathways. Most lines contain one missing *p* value, which is excluded from the occupancy sampling universe and conservatively assigned *p* = 1 for the 386-feature multiplicity audit.

At the reported *p* ≤ 0.10 threshold, low-fitness lines have 37, 60 and 39 flagged metabolites and occupy 21, 29 and 25 annotation bins. High-fitness lines have 25, 35 and 25 flags and occupy 17, 14 and 15 bins. The corresponding mean breadth values are 25.00 and 15.33. However, the mean number of flags differs by 17.00 metabolites. Selecting more metabolites also tends to occupy more bins, even when biological placement is exchangeable. We therefore compare each observed breadth with a null that preserves its hit count and observed category sizes (Fig. 1).

**Figure 1.**
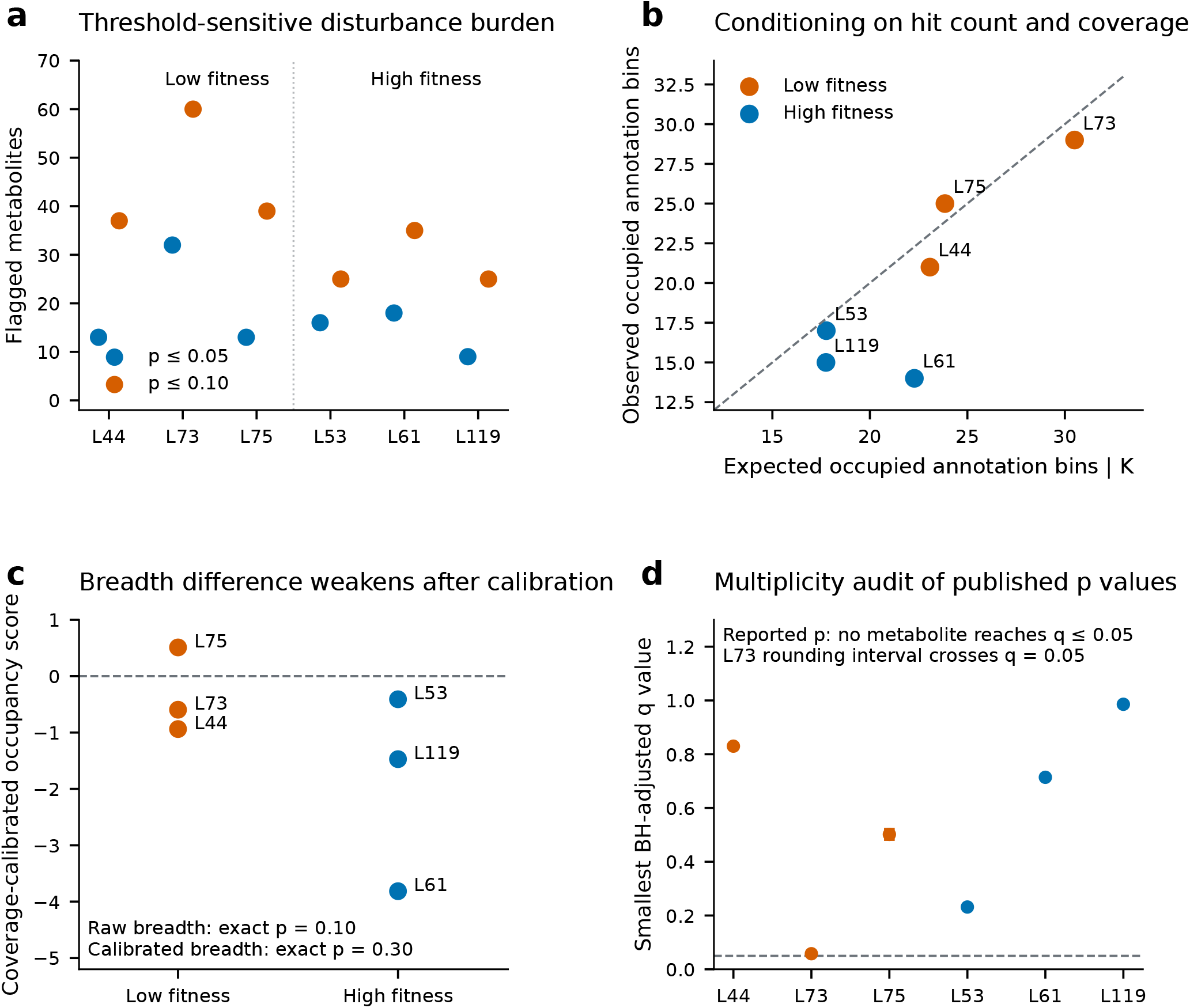
Coverage-conditioned audit of plant pathway disturbance. **a**, Counts of metabolites flagged at two reported significance thresholds. Each point represents one of six MA lines; metabolites are not independent biological replicates. **b**, Observed bin occupancy versus its exact conditional expectation at *p*≤ 0.10. The dashed line marks equality. **c**, Standardized occupancy residuals. Group comparisons enumerate all 20 three-versus-three label allocations. **d**, Smallest Benjamini–Hochberg-adjusted *q* values calculated from the reported *p* values, with intervals under a nearest-10^−4^ rounding assumption. L73’s interval crosses 0.05. Data are derived from the supplementary table of Kreutzmann et al.Kreutzmann et al. (2022).

Let *M* denote tested metabolites, *m*_*r*_ the size of disjoint annotation bin *r, K* the number flagged, and *S* the number of occupied bins. Under uniform sampling of *K* metabolites without replacement,

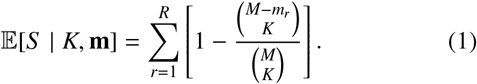

This is a coverage-conditioned reference, not a claim that metabolites have identical biological sensitivities. At *p* ≤ 0.10, expected breadth ranges from 17.75 to 30.50 bins. The two-sided exact group-label comparison yields *p* = 0.10 for raw breadth and *p* = 0.30 for the standardized occupancy residual. The direction of the group contrast remains positive; its evidential interpretation changes after accounting for the number of flagged features.

The calibrated statistic also reveals heterogeneity. L61 occupies 14 bins compared with an expectation of 22.28. Its exact conditional lower-tail probability is approximately 1.23 × 10^−4^ under the uniform-hit model, indicating unusually concentrated placement under that particular reference. This descriptive result does not establish a biological protection mechanism: metabolite correlations, unequal detectability and pathway-specific test power are not recoverable from the aggregate table. It does identify L61 as a candidate for a future replicate-level investigation of localized versus distributed responses.

The multiplicity audit requires similar care. Applying Benjamini–Hochberg correction (Benjamini and Hochberg, 1995) to the supplied *p* values yields no individual metabolite with *q* ≤ 0.05 in any line; the smallest reported-value *q* is 0.0579 in L73. Yet a plausible nearest-10^−4^ rounding interval moves this minimum between 0.04825 and 0.06755. A categorical claim that no metabolite could survive correction would therefore be stronger than the available precision supports. This audit concerns individual tests and does not invalidate an aggregate pathway association.

### 2.2. Metabolite-level patterns outperform disturbance burden in human diagnosis

We analyzed the public Z-score worksheet accompanying Thistlethwaite et al. (Thistlethwaite et al., 2022). It contains 539 profiles; 129 profiles enter our two explicitly separated benchmarks. The primary subset contains 40 EDTA profiles with argininemia (*n* = 13), argininosuccinic aciduria (*n* = 11), or ornithine transcarbamylase deficiency (*n* = 16). The secondary subset contains 89 heparin profiles covering nine diagnoses with at least five profiles per category. Keeping anticoagulant groups separate removes one direct source of cross-group discrimination but does not eliminate platform, cohort, age or treatment confounding.

All representations use the same repeated five-fold partitions. Feature inclusion and median imputation are estimated within training folds. A five-feature logistic burden baseline summarizes flag count, occupied-bin count, mean absolute deviation, root-mean-square deviation and maximum absolute deviation. Metabolite-level and pathway representations use a transparent cosine nearest-centroid classifier. These are fixed, interpretable benchmark configurations, not an optimized diagnostic system.

On the available panels, unsigned node profiles attain mean macro recall of 0.911 in the EDTA subset and 0.895 in the heparin subset. The burden baseline attains 0.624 and 0.382, respectively (Table 1; Fig. 2). The advantage of unsigned nodes over burden is 0.287 in EDTA and 0.513 in heparin. Conditional class-stratified sample-bootstrap intervals for these differences are 0.133–0.440 and 0.438–0.589. These intervals condition on the published profiles and validation predictions; they do not quantify performance at an external laboratory.

**Table 1.** Mean macro recall across 40 repetitions of five-fold internal validation. Repeated partitions reuse the same profiles and are not independent cohorts.

| Representation | EDTA | Heparin |
| --- | --- | --- |
|  | 3 diagnoses | 9 diagnoses |
| Burden only | 0.624 | 0.382 |
| Unsigned nodes | 0.911 | 0.895 |
| Unsigned graph | 0.909 | 0.895 |
| Unsigned pathway | 0.851 | 0.839 |
| Signed nodes | 0.908 | 0.867 |
| Signed graph | 0.904 | 0.856 |
| Signed pathway | 0.767 | 0.831 |

**Figure 2.**
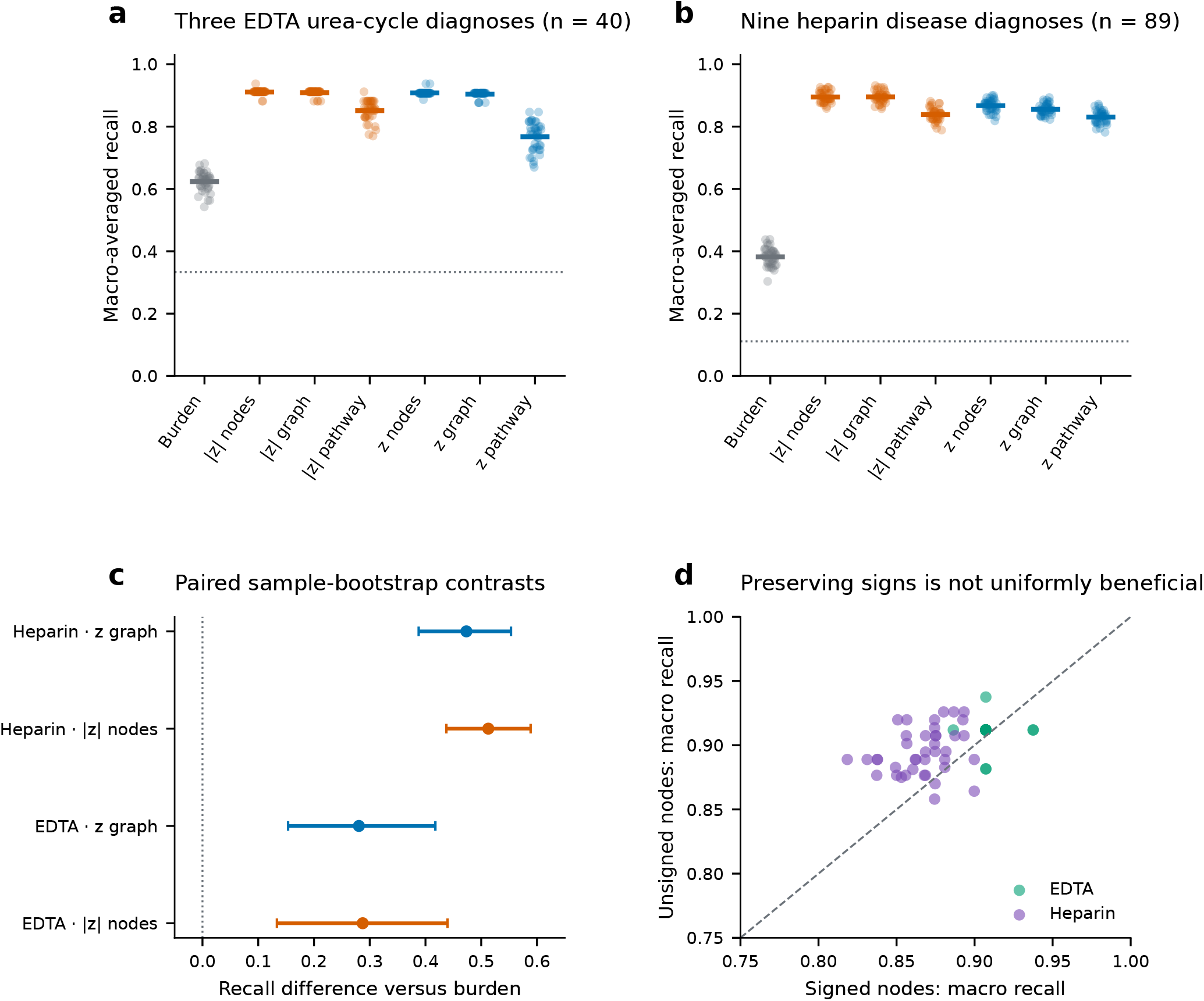
Diagnostic patterns retain information lost by scalar burden. **a,b**, Macro recall for seven representations in the EDTA and heparin subsets. Faint points denote 40 repeated validation partitions and horizontal bars their means. Dotted lines show reciprocal class count. **c**, Paired advantages over the five-feature burden baseline with 95% conditional class-stratified sample-bootstrap intervals. These intervals condition on the current profiles and predictions. **d**, Signed versus unsigned node performance for the same validation partitions. Neither points nor repetitions are independent external cohorts. Disease profiles are derived from Thistlethwaite et al.Thistlethwaite et al. (2022).

The comparison separates disturbance extent from disease identity. Different diseases can have similar overall burden while perturbing different metabolite coordinates. Nevertheless, the endpoint here is discrimination among selected known diagnoses. The results do not estimate sensitivity for detecting disease among healthy individuals, performance on unrepresented disorders, or clinical severity.

### 2.3 Pathway compression removes specific diagnostic contrasts

To assess information loss, we compare contrast retention and controlled measurement removal (Fig. 3).

**Figure 3.**
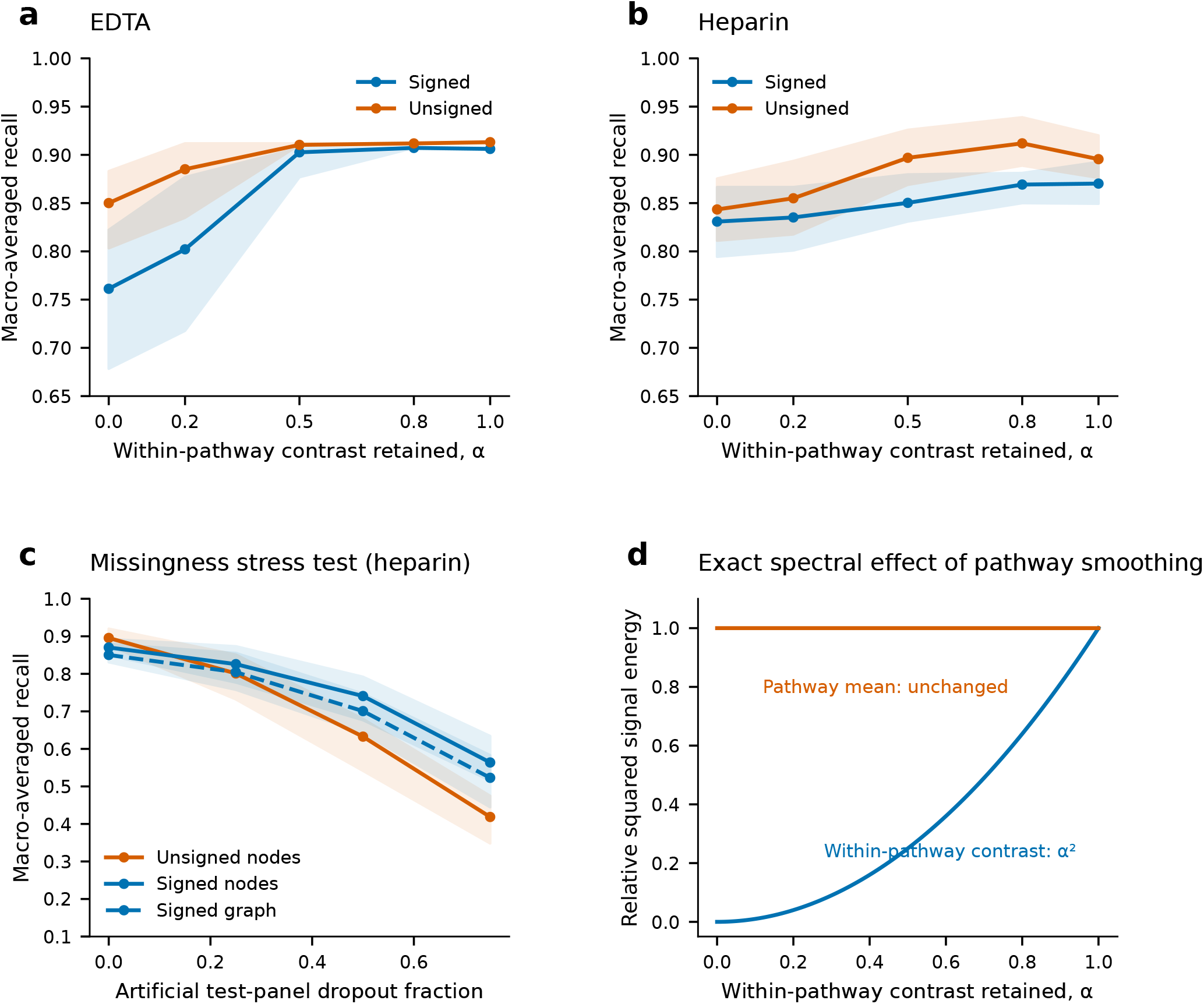
Information loss depends on the representation and assay coverage. **a,b**, Performance across the fraction of within-pathway contrast retained. At α = 0, all annotated features equal their pathway mean; at α = 1, node profiles are retained. Shading shows the 5th–95th percentile across 20 validation partitions, not an inferential confidence interval. **c**, Artificial removal of held-out measurements in heparin, with fixed training centroids and training-median imputation. Signed nodes become more robust than unsigned nodes as dropout increases. **d**, Exact squared-energy attenuation implied by the pathway projector. This panel is mathematical, while panels a–c are experimental-data analyses with artificial representation or coverage perturbations.

We construct a fixed co-membership graph from the supplied pathway labels. It is not a stoichiometric reaction network. Let *P* be the orthogonal projector that replaces each annotated feature by its pathway mean; unannotated features remain singleton nodes. The corresponding normalized within-bin Laplacian is *L* = *I P*. Graph regularization acts through the spectrum of its chosen operator (Smola and Kondor, 2003). For this co-membership construction, a graph-regularized profile is

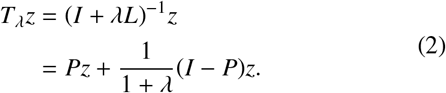

Thus smoothing preserves the pathway mean and shrinks within-pathway contrasts. Writing α = (1+λ)^−1^ makes the consequence explicit:

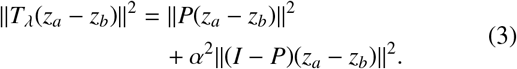

For two mean profiles separated only by a within-pathway contrast, complete averaging removes their separation exactly. This is a property of the projection, not a universal argument against network methods.

In the EDTA benchmark, signed pathway averaging lowers macro recall from 0.908 for signed nodes to 0.767. The paired conditional difference is 0.141, with interval 0.076–0.212. Moderate graph smoothing (*α* = 0.5) gives 0.904, retaining most of the metabolite-level information. In heparin, signed pathway recall is 0.831 versus 0.867 for signed nodes; its conditional difference interval includes zero. Our empirical result is therefore strongest in the primary subset. Neither graph representation demonstrates a general improvement over the corresponding node baseline on the available panels (Fig. 3).

### 2.4. Assay dropout reverses the ranking of signed and unsigned profiles

Available-panel performance slightly favors unsigned nodes, but preserving signs becomes advantageous when held-out measurements are artificially removed. In 20 repeated validation partitions, removing 50% of test features and imputing training medians yields mean macro recall of 0.741 for signed nodes and 0.633 for unsigned nodes in heparin. The signed-minus-unsigned difference is 0.108, with a conditional class-stratified sample-bootstrap interval of 0.063–0.155. At 75% removal the difference increases to 0.145 (interval 0.107–0.183). The corresponding EDTA effects are smaller and their intervals include zero.

The qualitative heparin reversal also appears under independent dropout seeds, shared omissions across each held-out panel, and zero rather than median imputation. At 50% dropout, signed-minus-unsigned node advantages in those additional configurations range from 0.084 to 0.140. The stress tests are controlled missingness experiments on measured profiles, not observations of a new assay. They indicate that representation ranking depends on coverage assumptions. They do not identify a clinically optimal panel or prove that preserving signs always helps.

There is a simple geometric reason why ranking can change. For feature-retention probability *q*, test profile *z*, training-imputation vector *m*, and normalized class centroids *c*_*a*_, *c*_*b*_, the expected signed classification margin under independent erasure is

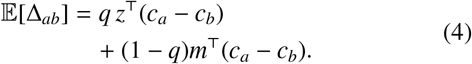

For unsigned profiles the analogous margin uses |*z*|, the unsigned centroids and the transformed imputation vector. The two margins need not decay at the same rate. Our results demonstrate the reversal in this dataset; Eq. (4) explains why completeness alone can change their ordering without any change in diagnosis.

### 2.5. Concentration normalization can conceal lower desired flux

The clinical benchmarks establish diagnostic information, not functional restoration. To make the distinction constructive, consider a metabolite pool *x* supplied at rate *u*, consumed by a desired product-forming route and by an alternative bypass:

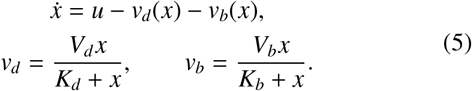

All parameters are positive. For equal affinities *K*_*d*_ = *K*_*b*_ = *K*, a finite stable steady state exists exactly when *u* < *V*_*d*_ + *V*_*b*_. Defining *V* = *V*_*d*_ + *V*_*b*_ and capacity reserve ϵ = *V* − *u* gives

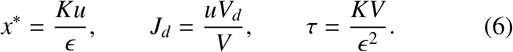

Here *τ* is the local relaxation time. Increasing bypass capacity decreases both concentration and desired product flux:

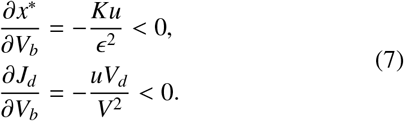

For *K* = 1, *u* = 0.8, *V*_*d*_ = 1, and *V*_*b*_ increasing from 0.1 to 2, concentration falls from 2.667 to 0.364 (86.4%), while desired flux falls from 0.727 to 0.267 (63.3%). Relaxation simultaneously accelerates: τ falls from 12.22 to 0.620. Both a smaller pool and faster recovery can therefore accompany lower function in this model (Fig. 4). Whether the trade-off is harmful depends on the actual task and any toxicity of the intermediate; the model is not a treatment recommendation.

**Figure 4.**
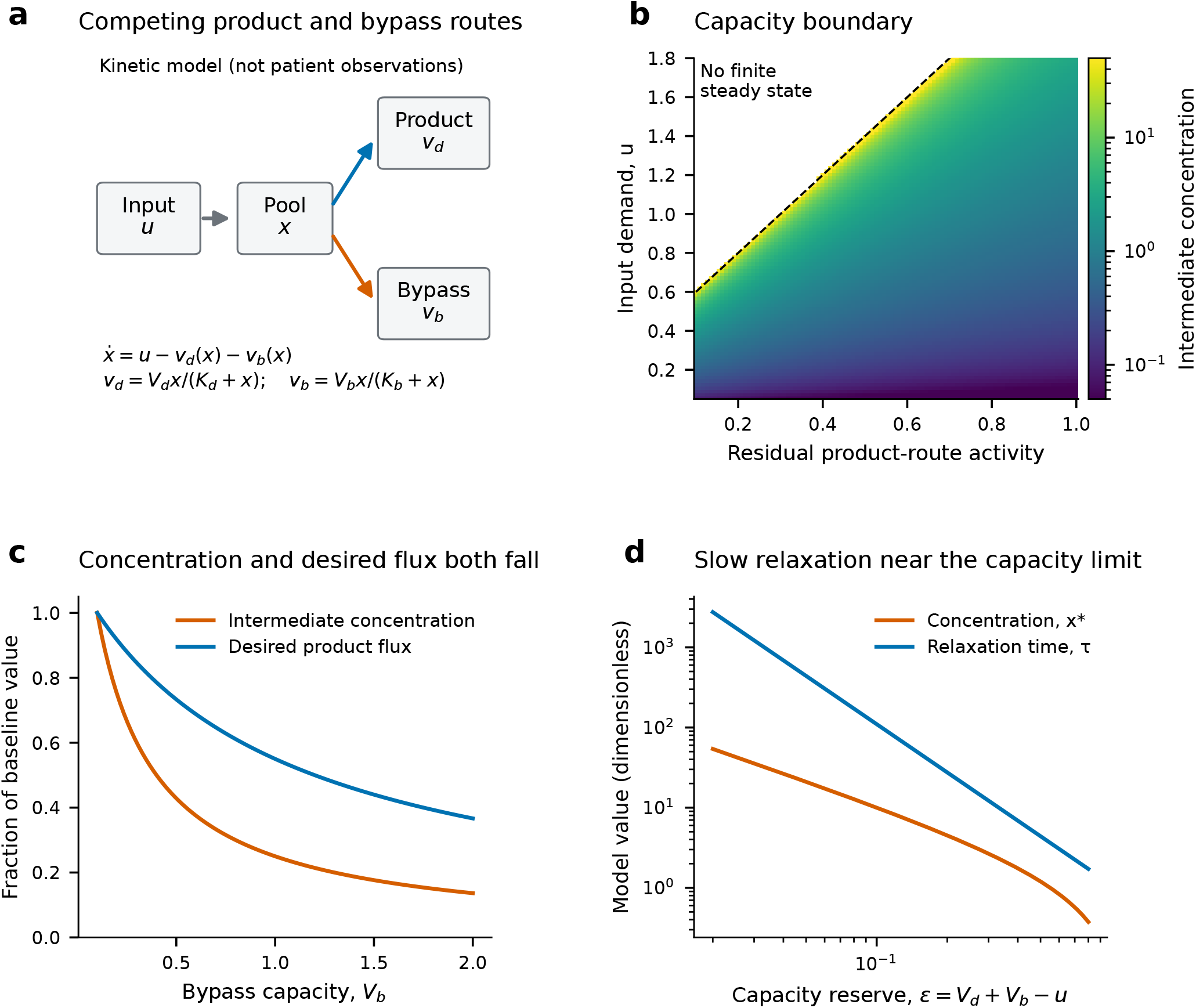
A capacity-constrained model separates observed pools from desired function. Every panel shows an uncalibrated kinetic model, not patient data. **a**, Input enters a shared intermediate with competing desired and bypass routes. **b**, Steady concentration across input demand and residual desired-route activity. The dashed boundary is *u* = 2*r* + 0.4; the white upper-left region has no finite steady state. **c**, Increasing bypass capacity reduces both the intermediate concentration and desired product flux, each normalized to its value at *V*_*b*_ = 0.1. **d**, As capacity reserve approaches zero, concentration and relaxation time increase at different rates. Equations (6)–(12) give exact solutions and identify what additional measurements can resolve.

## 3. A mathematical criterion for functional interpretation

Let a metabolic system satisfy 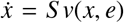, where *S* is the stoichiometric matrix and *e* is an enzyme-activity parameter vector. Work in independent state coordinates after eliminating any conserved pools. At a stable reference state with a nonsingular Jacobian, define *J = S ∂*_*x*_*v* and *B = S ∂*_*e*_*v*. For sufficiently small perturbation

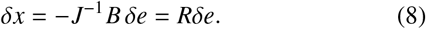

An assay observes *y = Hδx = Oδe*, with *O = HR*. A desired scalar flux response is *δ j = f* ^⊤^*δe*, where

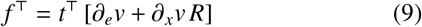

for a task weighting *t*. The metabolome locally identifies this task if and only if

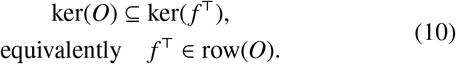

### Proposition 1

(Local task identifiability). *For known linear operators O and f* ^⊤^, *every pair of enzyme perturbations with the same observed response has the same task response if and only if Eq. (10) holds*.

*Proof*. Equal observations imply that the difference between the perturbations lies in ker(*O*). The task responses agree for every such difference exactly when *f* ^⊤^ annihilates this kernel. Since row(*O*) = ker(*O*)^⊥^, this is equivalent to the row-space condition.

This is standard linear identifiability applied to a metabolic endpoint (Villaverde et al., 2016). Failure permits identical observed metabolic changes with different task responses; scalar disturbance scores cannot resolve that ambiguity.

This observation also yields an exact local uncertainty bound. Suppose *y* is noise-free and compatible with ∥ *δe* ∥ ≤ ρ. Let *δe*_0_ = *O*^†^*y* be the minimum-norm solution, and let columns of *N* form an orthonormal basis for ker(*O*). Then the compatible task responses fill the interval

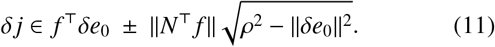

The bound is attained by choosing the null-space component parallel or antiparallel to *N*^⊤^ *f*; when *N*^⊤^ *f* = 0, the interval collapses to one value. If the interval crosses zero, the available measurements cannot certify the sign of the response under the stated model and perturbation bound. This interval is not fitted to our human profiles; the necessary patient-specific kinetic parameters are unavailable.

The kinetic example makes the measurement requirement particularly clear. Knowing *K, x*^*^ and τ identifies aggregate capacity and input:

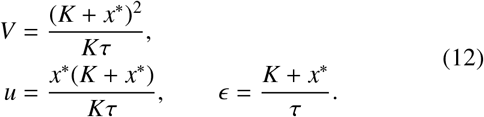

Yet these observations do not identify the split between *V*_*d*_ and *V*_*b*_. Pathway-specific flux information is still required to infer *J*_*d*_. Dynamic pool measurements can resolve aggregate reserve while leaving desired function ambiguous.

## 4. Discussion

The strongest empirical contribution is a coverage-dependent reversal in diagnostic representation ranking. On the available heparin panel, an unsigned profile is slightly more discriminative than a signed one; after controlled feature removal, signed profiles retain substantially more diagnostic information. This is a result about representation and measurement, not a newly discovered disease mechanism. Its reproducibility across omission structure and imputation choices makes external validation worthwhile, while the lack of an independent laboratory cohort prevents a clinical performance claim. Repeated partitions of a small collection cannot substitute for independent samples, and uncertainty across partitions is not external generalization uncertainty (Nadeau and Bengio, 2003; Varoquaux, 2018).

The plant reanalysis provides a complementary lesson. A broad disturbance count incorporates hit count, pathway sizes, significance thresholds and assay precision. Conditioning on these factors changes the strength of the low-versus-high fitness comparison. The concentrated response in L61 is an experimentally testable lead, but the uniform-hit reference does not account for within-pathway dependence. It should motivate targeted follow-up rather than a claim that localization causes higher fitness. Our analysis preserves the central value of the mutation accumulation design while sharpening what its metabolomics table can support.

The theoretical contribution separates diagnosis from function. A phenotype may be easy to classify yet insufficient to identify a desired metabolic task. Conversely, an intervention can reduce an abnormal intermediate while diverting material away from a necessary product. The explicit null-space criterion defines when an assay can support a functional interpretation; the kinetic example shows why even faster relaxation can be misleading. The distinction follows from established control and identifiability concepts (Kacser and Burns, 1995; Heinrich and Rapoport, 1974; Raue et al., 2009; Villaverde et al., 2016), applied here to an explicitly defined metabolic task.

For inherited metabolic disorders, the immediate research need is to combine interpretability with reliable measurement. A practical next study would validate representation rankings using paired measurements from different assay panels, with patient-level separation and treatment metadata. A mechanistic arm would measure targeted pool dynamics and pathway-specific isotope flux in a selected disorder, using isotope-constrained flux analysis to resolve routing (Zamboni et al., 2009). That design could test whether measured reserve and task flux improve interpretation beyond disease resemblance. It would also clarify whether changes in the same metabolite reflect a primary enzyme defect, treatment, diet or secondary compensation.

Network-based methods remain valuable when their edges and operators match the question. Methods for feature-to-network integration and gene prioritization (Li et al., 2013; Pirhaji et al., 2016; Graham Linck et al., 2020; Bongaerts et al., 2022) complement recent bipartite propagation work that expands topology-informed statistical signal beyond measured nodes (Kuligowski et al., 2026); it does not equate propagated signal with experimentally observed abundance. Our co-membership smoothing is simpler and should not be treated as a benchmark of every reaction-network algorithm. SBcoyote provides an extensible Python environment for reaction editing and SBML visualization (Xu et al., 2023); it is a suitable environment for inspecting the supplied kinetic model and future pathway-specific extensions. It was not used as a solver or to generate the present figures.

Several limitations delimit the interpretation. Six deliberately selected plant lines provide a narrow basis for population inference. Aggregate plant *p* values prevent replicate-level reanalysis, dependence-aware nulls and recovery of full test precision. Human Z scores inherit upstream reference normalization; fold-local processing cannot undo that history. Sample identifiers are unique, but independent patient identities across source studies cannot be reconstructed. Treatment, age, platform and ascertainment effects remain possible. Artificial random dropout does not represent every form of missingness, and no clinical severity, causal biomarker or intervention outcome is evaluated. Finally, the equal-affinity kinetic model is intentionally minimal and uncalibrated. Its counterexample is valid under its assumptions; its numerical values are not predictions for patients.

## 5. Conclusions

Assay coverage is part of the inference problem rather than a neutral property of the input. Conditioning on hit count and annotation-bin size changes the interpretation of pathway breadth, while controlled measurement removal can reverse the ranking of signed and unsigned diagnostic representations. Separately, the kinetic model and task-identifiability criterion show that observed pool improvement does not by itself identify functional recovery. Together, these analyses define testable requirements for robust metabolic diagnosis and mechanistic interpretation: validate representation choices across assay coverage, preserve task-relevant contrasts, and measure flux where concentration observations leave functional ambiguity.

## 6. Methods

### 6.1. Source data, provenance and analysis units

The plant input is publisher supplementary file `10682_2022_ 10210_MOESM1_ESM.xlsx`, sheet ST3, accompanying Kreutzmann et al. (Kreutzmann et al., 2022). ST1 supplies line fitness categories. The unit for between-group inference is an MA line, not an individual metabolite. ST3 supplies fold changes and *p* values but no replicate intensities. We calculate log_2_ fold changes without interpreting them as metabolic flux. The 59 category labels are retained verbatim; overlapping biochemical roles are not reconstructed from these bins. Annotation certainty is inherited from the source tables rather than established by new identification experiments; this distinction is consistent with metabolomics reporting standards (Sumner et al., 2007).

The human input is supplementary file `41598_2022_ 10415_MOESM2_ESM.xlsx`, Z-score worksheet, accompanying Thistlethwaite et al. (Thistlethwaite et al., 2022). Our parser recovers 539 profiles and 1,375 feature rows. The row count is a workbook parsing result, not a claim that all rows represent distinct identified metabolites. Unknown-diagnosis Alaimo profiles and reference profiles do not enter the known-disease classification task. Primary selection is restricted to EDTA and the three diagnoses listed above. Secondary selection is restricted to heparin diagnoses with at least five profiles: cerebral creatine deficiency syndrome 2 (8), citrullinemia (9), cobalamin biosynthesis (6), glutaric acidemia 1 (5), maple syrup urine disease (18), methylmalonic aciduria (9), ornithine transcarbamylase deficiency (17), phenylketonuria (8), and propionic acidemia (9).

Input files were retrieved on 3 October 2026 and hashed with SHA-256. Source URLs, hashes, derived tables, per-sample validation outputs and complete code are provided with the manuscript. Reproduction downloads the public publisher workbooks; this package does not redistribute the source workbooks. The reuse consists of analysis of public tables, not new sample collection.

### 6.2. Exact occupancy distribution and variance

Write 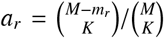 for the probability bin *r* receives no hit. The variance of *S* is

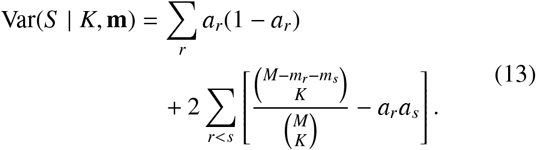

Combinations with fewer available elements than *K* are assigned zero. We report 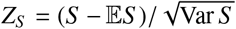 as a calibration statistic, without assuming its distribution is Gaussian. The full exact distribution follows from coefficient extraction:

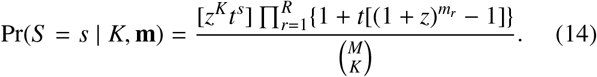

An iterative dynamic program accumulates coefficients for selected metabolite count and occupied-bin count. It preserves the category sizes exactly. Both moments and the full distribution are verified against enumeration of all 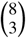 subsets for bins of sizes 2, 3 and 3. Lower-tail probabilities describe concentration under the uniform-hit reference; upper tails describe dispersion.

For group comparisons, all 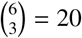 possible low/high label allocations are enumerated. The statistic is the difference between the means of the three low- and three high-fitness lines. The two-sided tail uses |*T* |≥ |*T*_obs_|, including the observed allocation. As complementary allocations have opposite statistics of equal magnitude, a nonzero two-sided exact *p* cannot be below 2/20 = 0.10. Exchangeability is a descriptive reference for these selected lines, not a randomized treatment assignment.

### 6.3. Threshold, multiplicity and magnitude summaries

Flags use the published *p* ≤0.05 and *p*≤ 0.10 thresholds. Bin occupancy requires at least one flagged metabolite. Benjamini–Hochberg adjustment is computed separately for each line across the 386 supplied features, with missing values set to 1. The rounding sensitivity assumes the displayed values arose by nearest-10^−4^ rounding and evaluates all *p* values at their lower and upper endpoints, clipped to [0, 1]. This is an assumption-dependent precision audit, not recovery of original test results.

Threshold-free summaries include RMS log_2_ fold change and an entropy-based effective number of energy-bearing bins, using the effective-number construction of Hill (1973),

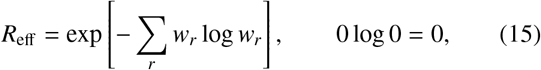

where *w*_*r*_ is the fraction of total squared log_2_ fold change contributed by bin *r*. These summaries describe effect magnitude and allocation but cannot supply missing biological replication.

### 6.4. Internal validation and classifier definitions

We use 40 repetitions of stratified five-fold validation with seed 20261003 + *r* for repetition *r*. Each held-out profile is excluded from centroid fitting, feature selection and median estimation. A feature enters a fold if it is observed in at least 80% of its training profiles; training medians fill existing missing values in training and test data. Values are clipped to [−10, 10] to limit the influence of extreme source Z scores, without selecting that threshold on test performance. The selected feature counts span 284–340 for EDTA and 477 for heparin; the authoritative ranges are also recorded in the output JSON.

The burden vector is

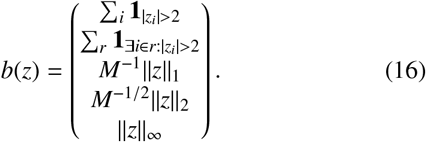

It is standardized using training means and variances and fitted with a class-balanced logistic regression with *C* = 1 and maximum 2,000 iterations. Feature-level representations use *z* or |*z*|. For class c, the training mean profile is normalized to unit Euclidean length, and a test profile is assigned to the centroid with the largest dot product. This is equivalent to cosine nearest-centroid classification for nonzero test profiles.

Graph profiles use Eq. (2) with λ = 1, giving α = 0.5; pathway profiles use *Pz* or *P z*. Unannotated rows remain independent singletons. The contrast-retention experiment uses α 1, 0.8, 0.5, 0.2, 0 across 20 repetitions. These parameters are fixed analysis choices. We do not select a best parameter and then report its test performance. The present benchmark is a comparison of representations and deliberately simple modeling choices, not a controlled comparison of all possible learners.

Macro recall is the mean class-specific recall and gives equal weight to diagnoses. Ordinary accuracy is also provided in the result tables. Split-level scatter and shaded percentile bands show variability across partitions of the same data. They are not independent sampling distributions.

### 6.5. Dropout, robustness and conditional uncertainty

The principal dropout experiment uses 20 validation repetitions and independent Bernoulli removal of each test feature at fractions 0, 0.25, 0.50 and 0.75. Removed values are replaced with the corresponding training median, while the trained centroids and feature selections remain fixed. Dropout is additional to the source dataset’s existing missingness. Its random seeds depend only on the fixed seed, repetition, fold and dropout fraction.

Additional configurations use ten repetitions, independent dropout seeds, either individual-profile masks or a mask shared by all test profiles in a fold, and either median or zero imputation. We also provide a global sign-reversal stress test that preserves absolute magnitude while changing the sign of a random half of test profiles. That test characterizes fragility to a deliberately artificial transformation; it is not a model of disease biology.

We use a conditional, class-stratified application of bootstrap resampling (Efron, 1979). For paired conditional uncertainty, we first average each sample’s correctness across repetitions, calculate between-model differences, then resample profiles with replacement within each diagnosis 3,000 times. The reported percentile interval covers the resulting mean class-balanced difference. Repeated cross-validation predictions share training data, and this conditional bootstrap does not incorporate all training instability or external-site heterogeneity (Nadeau and Bengio, 2003; Varoquaux, 2018). No *p* value is assigned to the coverage crossover, and its small-dataset status remains explicit.

### 6.4 Kinetic calculations and verification

For Fig. 4, all variables are nondimensionalized. The phase grid spans *u* = 0.05–1.8 and residual desired-route activity *r* = 0.1–1, with *V*_*d*_ = 2*r, V*_*b*_ = 0.4 and *K*_*d*_ = *K*_*b*_ = 1. No finite steady state is reported where *u* ≥ *V*_*d*_ + *V*_*b*_. Elsewhere the steady state is independently calculated by bracketed root finding and checked against mass balance. The intervention comparison fixes *u* = 0.8 and *V*_*d*_ = 1 and varies *V*_*b*_ = 0.1–2. The code verifies the negative concentration and product-flux derivatives numerically.

The relaxation expression is derived from 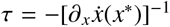. Stability holds because 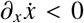. For the general linear criterion, assume a differentiable rate law, a locally invertible stable Jacobian, known observation operator and a sufficiently small perturbation. Eq. (11) follows by decomposing every compatible perturbation as *O*^†^*y* + *Nw* and maximizing *f* ^⊤^*Nw* over its Euclidean ball. It is an exact bound for the linear model; measurement noise, parameter uncertainty and nonlinear departures would enlarge or modify the compatible set.

### 6.7 Software and reaction-model exchange

Analyses use Python 3.12.14, NumPy 2.3.5, pandas 2.2.3, SciPy 1.17.0 (Virtanen et al., 2020) and scikit-learn 1.8.0 (Pedregosa et al., 2011); plotting uses Matplotlib 3.10.8 (Hunter, 2007). Figures are supplied as vector PDF/SVG and 600 dpi PNG with a color-accessible palette. The kinetic model is provided in SBML (Hucka et al., 2003) and Antimony (Smith et al., 2009) for reaction-level inspection and extension; its SBML consistency check reports no errors. SBcoyote (Xu et al., 2023) is cited for its Python-based editing, visualization and plugin architecture; its GUI was not executed in this analysis. We do not claim model fitting, SBcoyote-generated plots or patient-specific kinetic calibration.

## 7. Data and code availability

The accompanying reproducibility package contains the analysis scripts, per-line and per-profile outputs, conditional uncertainty estimates, figure sources, SBML/Antimony model and input-file hashes. Experimental inputs are available from the publisher supplements of Kreutzmann et al. (2022) and Thistlethwaite et al. (2022); the download script records source URLs and SHA-256 provenance. The analysis uses only public supplementary tables and performs no new sample collection.

## Declaration of AI-assisted preparation

AI-assisted tools were used to help prepare the manuscript text and analysis code. The empirical calculations were performed in Python using the cited public supplementary tables, and the scripts and numerical outputs are supplied in the reproducibility package.

## Notes

### Competing Interest Statement

The authors have declared no competing interest.

